# Characterization of Cancer-Reactive T Cells and Neoantigen-specific T Cell Receptors

**DOI:** 10.1101/2025.10.29.685473

**Authors:** Wei Sun, Si Liu, Bo Yu, Zhehao Zhang, Philip Bradley, Marie Bleakley

## Abstract

Not all tumor-infiltrating T cells are cancer-reactive (CR). Several studies have reported gene expression signatures associated with CR-T cells. To integrate these findings, we developed a computational workflow, CAT (Cancer-Associated T cells), which harmonizes existing CR-T cell signatures and applies them to identify CR-T cells across an atlas of one million T cells. Our findings reveal that the abundance of CR-T cells varies across cancer types and that baseline levels of CR-T cells predict patients’ responses to immunotherapy. In parallel, we established a high-throughput computational platform, Neo-TCR, for systematic screening of neoantigen-specific TCRs and their cognate neoantigens. The efficacy of Neo-TCR is validated by cross-validation studies and replications in two independent datasets. Together, our findings suggest a new direction for developing biomarkers for cancer detection and monitoring: integrating CR gene expression signatures with neoantigen-specific TCRs.

## Introduction

T cells have the remarkable ability to recognize tumor cells by detecting tumor-specific antigens, often in the form of neoantigens, which are typically peptides generated as a result of somatic mutations. Not all tumor-infiltrating T cells are cancer-reactive (CR); many are bystanders that do not react to tumor cells [1]. For example, Oliveira et al. [2] reported that 4.7–43.9% of tumor infiltrating CD8+ T cells were CR. Recent studies have demonstrated that CR T cells exhibit distinct gene expression signatures [2–7]. Typically, these studies first identified a limited number of CR T cells, then leveraged single-cell RNA and TCR sequencing to trace more T cells from the same clones and characterize their transcriptional profiles. This approach has yielded gene sets with differential expression in CR T cells, referred to as gene signatures. While these pioneering works have advanced the field, each has primarily focused on the comparative advantage of its own signatures within its own dataset. Harmonizing these signatures will enable broader and more systematic investigations of CR T cells in future studies.

An important research direction closely related to the detection of CR T cells is the identification of cancer neoantigens and their corresponding neoantigen-specific TCRs. These efforts are essential for translational applications, such as the development of personalized cancer vaccines and TCR-engineered T cell therapies. Multiple experimental strategies have been developed to address this challenge; however, despite their effectiveness, these approaches remain labor-intensive, technically demanding, and time-consuming.

In this study, we developed two complementary computational platforms to investigate CR-T cells and neoantigen-specific TCRs. The first platform, CAT (Cancer-Associated T cells), integrates multiple CR signatures with additional filtering steps, including a neural network classifier, to robustly identify CR-T cells. We assembled a comprehensive atlas comprising one million T cells derived from tumors and matched adjacent normal tissues or peripheral blood. Application of CAT enabled the identification of approximately 70,000 CR-T cells, accompanied by detailed annotations of their gene expression signature scores and corresponding TCR sequences. We also demonstrated that the abundance of CR-T cells at baseline can predict immunotherapy response.

The second platform, Neo-TCR, provides an efficient computational framework for high-throughput screening of public neoantigens and their corresponding TCRs. Unlike most existing studies that focus primarily on personalized neoantigens, Neo-TCR prioritizes public neoantigens shared across patients, thereby informing the development of broadly applicable, cost-effective, off-the-shelf therapeutic products. Using this approach, we identified 47 somatic mutations likely to encode neoantigens, eight of which have previously been reported to generate immunogenic neoepitopes, providing proof-of-principle validation of our methodology. Moreover, combining the resources from CAT and Neo-TCR, we found that T cells carrying neoantigen-associated TCRs exhibit stronger CR signatures. This observation suggests that integrating CR gene expression signatures with neoantigen-specific TCRs can enhance both the sensitivity and specificity of identifying T cell–based biomarkers for cancer detection and monitoring.

## Results

### Transcriptomic characterization of CR T cells

We collected gene expression signatures of CR T cells from six previous studies [2–5, 8, 9], including positive signatures (genes upregulated in CR T cells) and negative signatures (genes downregulated in CR T cells or upregulated in virus-responding T cells). In total, we assembled five positive and two negative signatures for CD8+ T cells, and four positive and two negative signatures for CD4+ T cells (Methods and Supplementary Table 1).

We studied these gene expression signatures in ∼ 500,000 CD8+ T cells and 500,000 CD4+ T cells from five studies [10–14], which covered 383 cancer patients of 11 cancer types (Figure 1(A)). The transcriptomic count data from these one million T cells were processed by the same workflow for read-depth correction and quality controls. We scored each gene signature by single cell gene set enrichment analysis (scGSEA) if the gene set contained more than 30 genes, or by average gene expression otherwise. Then for each signature, we scaled all its scores across all CD8+ T or CD4+ T cells to have mean 0 and standard deviation of 1. The scores of different signatures are highly consistent (Figure 1(B), Supplementary Figure 1(A)). We then averaged all positive or negative signature scores to obtain a positive or negative score for each cell, respectively. As expected, T cells from tumor tend to have higher positive scores than T cells from adjacent normal or blood (Figure 1(C), Supplementary Figure 1 (B)).

**Figure 1:**
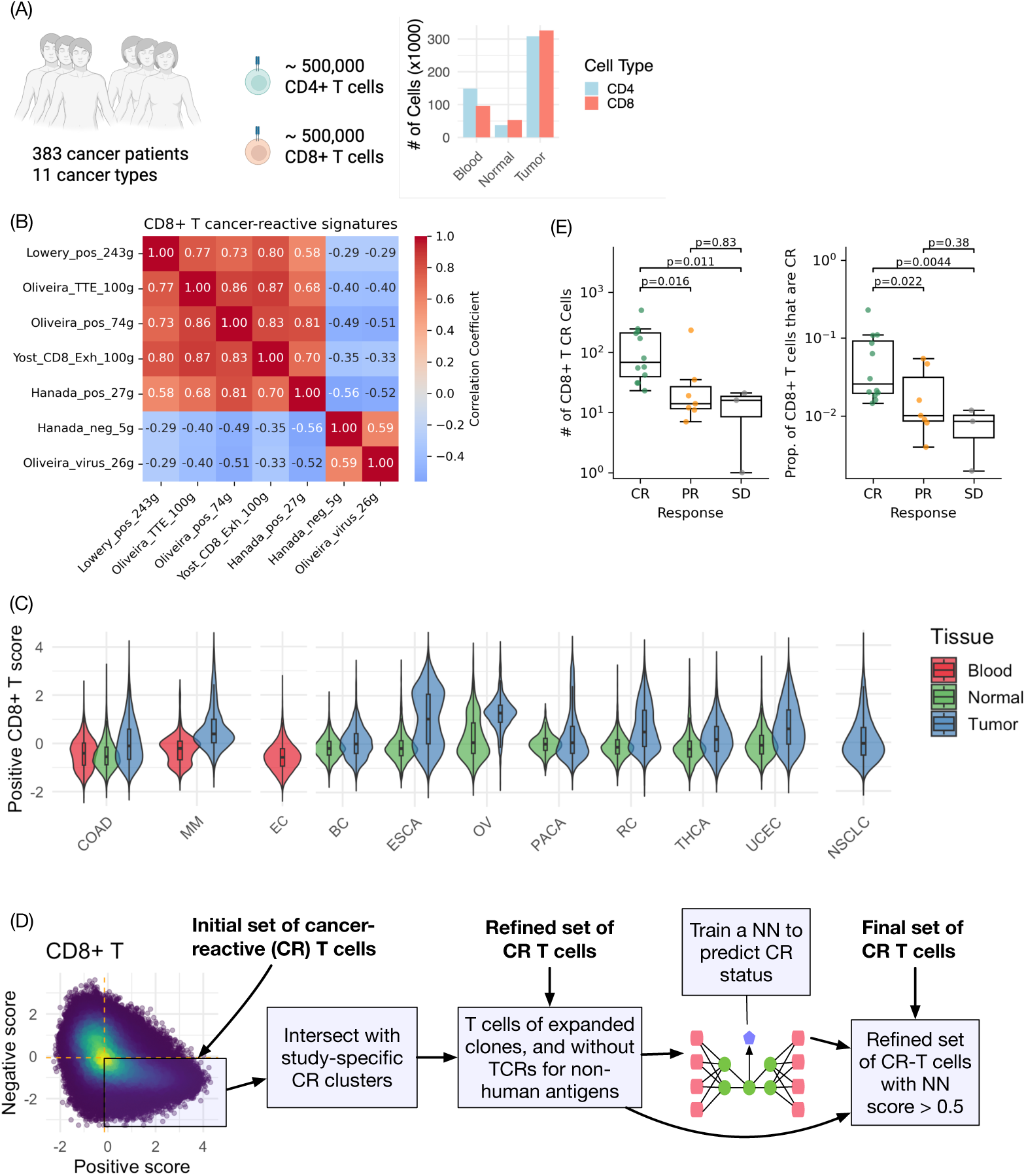
**(A)** An atlas of T cells collected from five studies. **(B)** Each CD8+ T cell is scored by the expression of seven signatures (gene sets). This heatmap showed the correlation matrix of those scores. Two gene sets with low expression in CR CD8+ T cells are labeled with “neg” or “virus” in their names. The other five gene sets are genes with high expression in CR CD8+ T cells. **(C)** A composite CD8+ T cell positive score was calculated by averaging the five individual signatures. This plot illustrates the distribution of this score across tissue types and cancer types. A corresponding analysis for CD4+ T cells, including the full names of the cancer types, is presented in Supplementary Figure 1(A). **(D)** A four-step workflow to identify a subset of high-confidence CR T cells. **(E)** Both the total number5of CD8+ CR T cells and the proportion of them among all CD8+ T cells, measured at baseline before treatment, are significantly associated with clinical outcome of immunotherapy of colorectal cancer [13].

Next, we annotated a T cell as CR or not by a four-step workflow (Figure 1(D), Supplementary Figure 2). First, an initial set of CR T cells was selected as those with higher positive scores and low negative scores (using the median of each score, respectively). In parallel, for each study, we clustered CD8+ and CD4+ T cells using the expression of the genes involved in the CR signatures, and identified CR clusters as those enriched with T cells having high positive scores and low negative scores. As the second step of the workflow, we intersected the initial set of CR T cells with study-specific CR clusters. In the third step, we selected T cells whose TCRs were observed in at least three cells in the whole dataset, which could be due to clonal expansion (i.e., three cells in one individual) or TCR sharing across individuals. We also removed a T cell from the CR set if its TCRs were annotated to be associated with non-human antigens, according to VDJdb [15] (Supplementary Figure 3). Finally, using the refined labels of CR T cells from step 3, we trained a neural network to predict cancer reactive status using genome-wide transcriptomic data (Supplementary Figure 4, and more details in the Methods Section). The prediction score ranges from 0 to 1. A T cell from the refined set was annotated as CR if the prediction score was larger than 0.5. The final set of CR CD8+ and CD4+ T cells include 62,188 and 7,322 cells, respectively. The relatively low number of high-confidence CD4+ CR T cells is likely due to the relatively lower consistency between CD4 CR signatures (Supplementary Figure 1(A)).

To demonstrate the use of CAT resources, we evaluated the association between the abundance of CD8+ T CR cells at baseline (before checkpoint immunotherapy) and the response to treatment in colorectal cancer [13]. Either the total number of CD8+ T CR cells or the proportion of them among all CD8+ T cells showed clear pattern that higher abundance of CD8+ T CR cells was associated with better response (Figure 1(E)).

### Association between TCRs and somatic mutations

The availability of CAT resources allows us to test a hypothesis: whether we can distinguish CR vs. non-CR T cells based on their TCRs. We fit multiple neural networks to assess this hypothesis and found that there was low accuracy in distinguishing CR TCRs from non-CR TCRs (Supplementary Figures 5, 6). This is likely due to the heterogeneity of CR TCRs. Previous studies have shown that epitope-specific TCRs tend to share similar sequences [16, 17]. However, given the large number of somatic mutations that could potentially give rise to neoantigens and high polymorphism of HLA genes in the human population, CR TCRs may need highly diverse sequences to recognize very different targets. Instead of identifying CR TCRs by their sequences, we next sought to prioritize neoantigen-specific TCRs by association analysis between TCRs and somatic mutations.

We collected TCR and somatic mutation data from over 8,000 cancer patients using data from The Cancer Genome Atlas (TCGA) [18]. TCR sequences were reconstructed from RNA-seq data of bulk tumor samples. As expected, the number of TCRs per sample was correlated with the estimated abundance of tumor-infiltrating T cells (Supplementary Figure 7). In total, we obtained 140,056 TCR alpha chains and 192,002 TCR beta chains. The vast majority of these were unique to individual patients and therefore are unsuitable for association analysis. We restricted our analysis to 8,463 patients with both somatic mutation data and TCR data. After filtering out somatic mutations and TCR chains that appeared in fewer than five patients, we ended up with 4,619 somatic mutations, 2,048 alpha chains and 867 beta chains. Associations between each somatic mutation and each TCR chain were assessed by a logistic regression that accounts for TCR read-depth and possible interaction between TCR read-depth and somatic mutation (Supplementary Figure 8, see Method Section for details). We also confirmed that the association results are not confounded by cancer types (Supplementary Figures 9-10, see Method Section for details).

### Somatic mutations status can be predicted using TCRs

Next, we sought to predict the occurrence of somatic mutations using their associated TCRs (Figure 2(A)). To obtain un-biased estimates of prediction accuracy, we implemented a cross-validation framework using TCGA data. Specifically, for 25 cancer types with at least 100 patients each, we trained mutation-specific predictors using 24 cancer types and evaluated their performance in the held-out cancer type. For each mutation, the predictor was defined as the number of TCRs associated with that mutation in the training data. Somatic mutations that can be predicted by TCRs are more likely to harbor public neoantigens than other somatic mutations, and subsequent analysis of peptide presentation by HLAs can prioritize neoantigens [20, 21] (Figure 2(A)).

**Figure 2:**
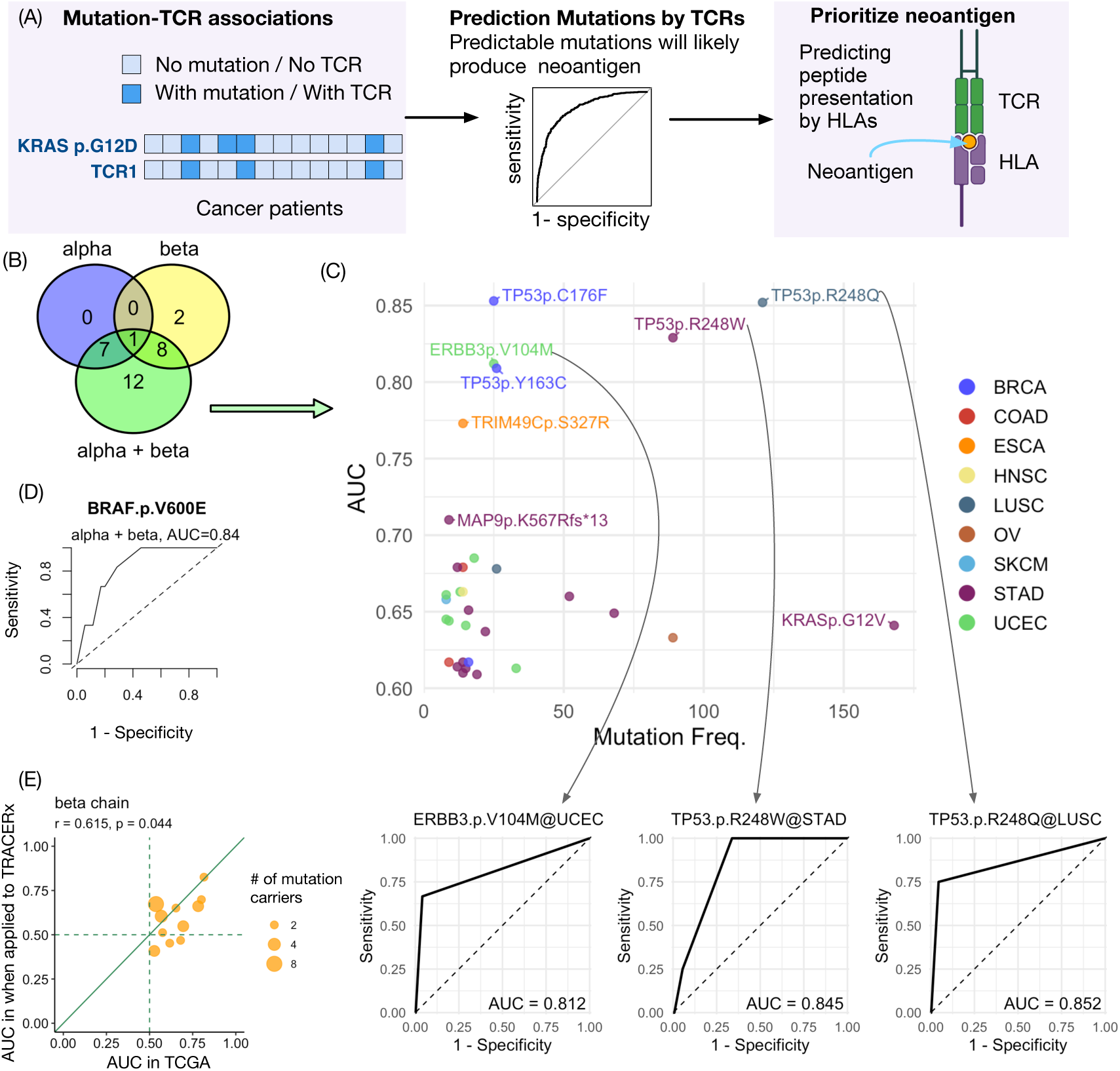
**(A)** An overview of the procedure to prioritize neoantigens by three steps: identify mutation-associated TCRs, predict mutation by TCRs, and prioritize neoantigens as peptides around the predicable mutations. **(B)** The number of predictable mutations with AUC high enough so that (1) AUC *>* 0.6 and (2) the corresponding FDR for AUC is smaller than 0.2. **(C)** A scatter plot of predictable mutations’ mutation frequency versus their AUCs, together with ROC curves for three examples. **(D)** Validation of the predictive model for BRAF V600E mutation using the scTCR data and somatic mutation data from Zheng et al. [10], using both alpha and beta chains with p-value cutoffs 0.1 for alpha chain and 0.2 for beta chain. Additional results in Supplementary Figure 11. **(E)** Validation of AUC predictions using the TRACERx data [19], using beta chain only with p-value cutoff 0.1. Additional results in Supplementary Figure 12.

In the TCGA dataset, TCR alpha and beta chains were not paired. Therefore, predictions were first generated separately for each chain and then combined. For each left-out cancer type, we considered somatic mutations present in at least three patients and assessed prediction performance using the area under the receiver operating characteristic (ROC) curve (AUC). Because we tested many mutations across 25 cancer types, some AUCs could appear high by chance. To account for this, for each p-value cutoff, we retained the predictions whose AUCs were high enough so that the corresponding FDRs were smaller than 0.2 (Supplementary Figures 11 and 12). The p-value cutoff was treated as a hyper-parameter. For the prediction using alpha and beta chains alone, we used p-value cutoff 0.1. For the predictions using both alpha and beta chains, which will be used in downstream validations, we merged the results of three sets of (TCR alpha, TCR beta) p-value cutoffs: (0.01, 0.05), (0.05, 0.1), and (0.1, 0.2) to provide a broader set of candidates for downstream validation. Here a relatively smaller p-value cutoff was chosen for alpha chain because alpha chains tend to have less diversity and higher population frequencies, and thus TCR-mutation associations tend to have smaller p-values in association testing.

Overall we could predict 30 mutations with AUC high enough (FDR *<* 0.2, and AUC *>* 0.6) in the combined results of alpha chain only, beta chain only, and alpha + beta chains, while 28 out of these 30 mutations were captured when using both alpha and beta chains (Figure 2(B)) and six of these 28 mutations can be predicted with AUC *>* 0.7 in different cancer types (Figure 2(C)).

Next, we trained three prediction models for each somatic mutation using the full TCGA dataset, applying three pairs of p-value thresholds for TCR alpha and beta chains, from stringent to liberal: (0.01, 0.05), (0.05, 0.1), and (0.1, 0.2). The number of mutations that are predictable (i.e., those with at least one associated TCR) are 4,600, 4,607, and 4,610, respectively.

We then evaluated these models in two independent datasets. The first dataset consisted of somatic mutation and single-cell TCR (scTCR) data from 41 cancer patients reported by Zheng et al. [10] (Figure 2D). Among these patients, only one mutation occurred in more than three individuals—the BRAF V600E mutation, present in 6 of the 41 patients. The models trained on TCGA data accurately predicted BRAF V600E status in this cohort, with the best performance achieved using both alpha and beta chains under the most liberal cutoff pair (0.1, 0.2) (Figure 2B). Notably, across all three cutoff settings, the models yielded consistently high performance, with AUCs exceeding 0.7 by using beta chain only or both alpha and beta chains (Supplementary Figure 13).

The second validation dataset was obtained from the lung TRACERx (Tracking Cancer Evolution through Therapy) study [19], which included TCR alpha and beta chain sequences as well as somatic mutation profiles from 65 lung cancer patients. Due to the limited sample size, most somatic mutations were private to individual patients, precluding a systematic evaluation of prediction accuracy. Among the somatic mutations for which prediction models had been trained on TCGA data, 11 were observed in at least two TRACERx patients. When applying the TCGA-trained models for these 11 mutations, predictions based on TCR beta chains showed strong concordance between TCGA and TRACERx, whereas predictions based on TCR alpha chains showed no reproducibility (Figure 2E, Supplementary Figure 14). The greater consistency of beta chain–based predictions may reflect the higher mutation specificity of TCR beta chains; however, given the limited sample size, these findings should be interpreted with caution.

### T cells with mutation-associated TCRs tend to have CR gene expression signatures

We have compiled CAT, a T cell atlas with positive/negative CR scores as well as CR labels (Figure 1D). Here, we evaluated whether T cells with mutation-associated TCRs have CR gene expression signatures. We classified T cells in CAT into four groups by matching their TCRs with mutation-associated TCRs: matching of neither alpha or beta chain, matching by alpha or beta alone, and matching both chains. Interestingly, if both the alpha and beta chains of a CD8+ T cell are mutation-associated TCRs, it is three times more likely to be classified as a CR CD8+ T cell (50% vs. 17% for neither, Figure 3(A)). In contrast, if both the alpha and beta chains of a CD4+ T cell are mutation-associated TCRs, its chance to be classified as a CR CD4+ T cell is only slightly higher (2.2% vs. 1.9% for neither, Figure 3(D)). Such enrichment of CR signatures was not observed for T cells that matched by alpha or beta chain only. Positive/negative scores derived from gene expression signatures strongly support that matching of both TCR chains increases the likelihood of being CR (Figure 3(B-C)). The pattern of positive/negative scores are less clear for CD4+ T cells, likely due to less informative gene expression signatures.

**Figure 3:**
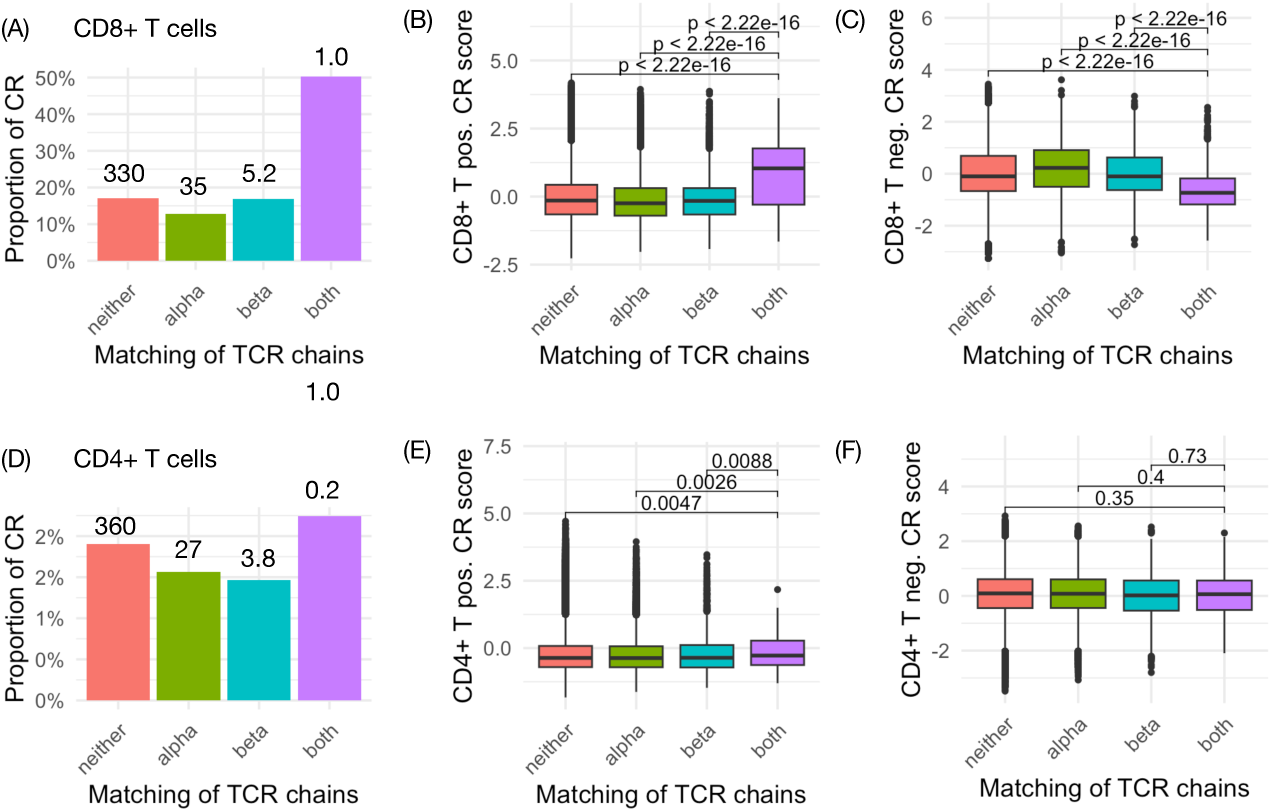
(A,D) The proportion of cells being cancer-reactive for CD8+ T cells and CD4+ T cells across four groups of cells, classified by matching their TCRs with mutation-associated TCRs identified from the TCGA data. The groups are defined as follows: alpha chain only (alpha), beta chain only (beta), both chains (both), or neither chain (No). The number on top of each bar indicates the number of cells (by 1000). (B-C) Positive and negative CR scores for CD8+ T cells. (E-F) Positive and negative CR scores for CD4+ T cells (C-D)

### Characterization of TCR-associated somatic mutations

TCR-associated somatic mutations are more likely to encode public neoantigens, which represent promising targets for immunotherapy and cancer vaccine development [22]. To systematically characterize these mutations, we analyzed 5,223 *public* somatic mutations that appear in at least five cancer patients and classified them into three categories based on their strongest statistical association with any TCR. Specifically, we designated mutations as (1) non-immunogenic (“No”) if their minimum association p-value (*p*_min_) is larger than 0.1 (1,265 mutations), (2) moderately associated (“Moderate”) if 0.001 *< p*_min_ ≤ 0.01 (1,908 mutations), and (3) strongly associated (“Strong”) if *p*_min_ ≤ 0.001 (2,050 mutations) (Figure 4A).

**Figure 4:**
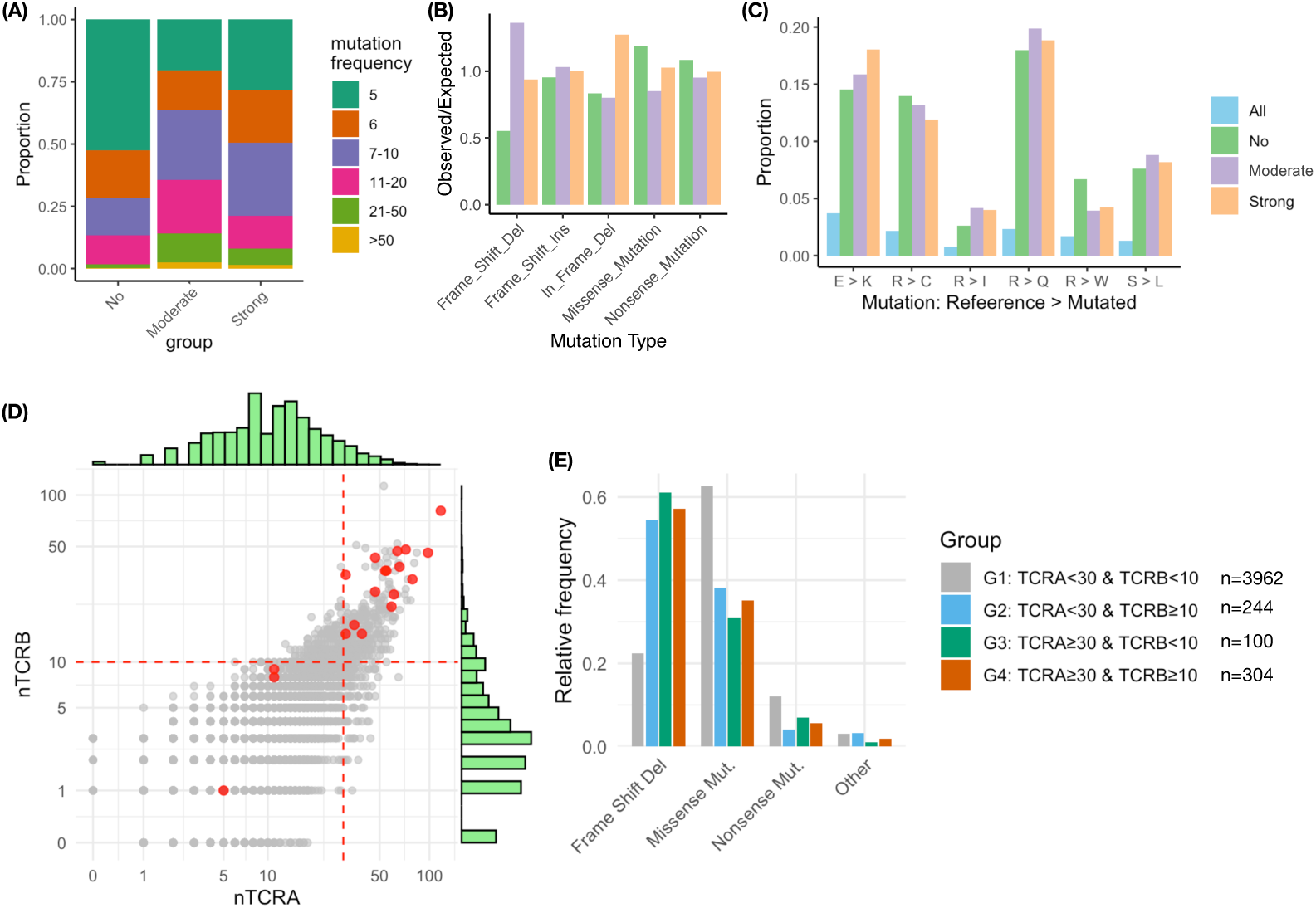
**(A)** Summary of the mutation frequencies for three groups of public somatic mutations that appear in at least five patients: No (not associated with any TCR), Moderate (with moderate association 0.001 *< p* ≤ 0.1 for at least one TCR), and Strong (strongly associated with at least one TCR with *p <* 0.001). **(B)** Enrichment or depletion of different types of somatic mutations compared to what is expected by chance. **(C)** Frequencies of different types of missense mutations. **(D)** A scatter plot of somatic mutations by the number of associated TCR alpha and beta chains. **(E)** The relative frequencies of different groups of somatic mutations (grouped by the number of associated TCRs) stratified by the type of mutations.

For frame-shift deletions, there is a relative depletion of non-immunogenic mutations (group No, Figure 4 (B), Supplementary Table 3). This is consistent with previous findings that indels tend to generate more immunogenic peptides [23]. Among missense mutations, the frequencies of different substitution types were comparable across the three groups of public mutations, but there are substantial differences between these public mutations and all mutations (Figure 4 (C)), suggesting a systematic difference between public mutations and rare or private mutations.

Somatic mutations exhibiting moderate versus strong associations with individual TCRs did not display substantial differences. This likely reflects the limited reliability of the association signals between a somatic mutation and a single TCR, due to data sparsity. To overcome this limitation, we instead characterized each somatic mutation by the number of distinct TCRs with which it was significantly associated (Figure 4D). We further curated a set of somatic mutations with experimentally validated neoantigens by integrating data from NEPdb [24] and several studies [25–32]. Notably, mutations with known neoantigens tended to be associated with a larger number of TCRs (Figure 4D), and those mutations associated with multiple TCRs were enriched for frameshift deletions (Figure 4E). Collectively, these findings suggest that the breadth of TCR associations, instead of the strongest association, provides a more robust criterion for prioritizing somatic mutations likely to encode neoantigens.

Therefore, we selected the 25 somatic mutations that were associated with the most TCRs (more than 50 TCR alpha chains and more than 30 TCR beta chains) as potential candidates that encode neoantigens. Combined with the 28 somatic mutations prioritized by the cross-validation study, we ended up with 47 unique somatic mutations (Supplementary Table 5). Among them, eight are known to encode neoantigens that can induce immune responses (Supplementary Table 6). The remaining 39 of them are potential candidates for future studies to identify novel public neoantigens.

## Discussion

We have developed two complementary approaches to identify biomarkers for cancer detection and monitoring. CAT (Cancer-Associated T cells) detects cancer-reactive T cells based on their gene expression signatures, while NeoTCR identifies neoantigen-specific TCRs through association analysis. Although several studies have previously reported cancer-reactive gene expression signatures, our contribution lies in assembling and aggregating these signatures within a unified workflow with additional filtering and validation steps. The feasibility of identifying neoantigen-specific TCRs by association analysis is supported by analogous studies in which TCR repertoires accurately predicted viral antigen exposure [33,34]. Although one TCR can recognize many antigens and one antigen can be recognized by multiple TCRs [35], we expect aggregating multiple TCRs that are associated with a somatic mutation can yield accurate predictions of mutation status. Our results also demonstrate the convergence of these two approaches, i.e., the T cells with mutation-associated TCRs are more likely to have cancer reactive gene expression signatures.

We demonstrate that the proportion of CR T cells prior to immune check-point inhibitor (ICI) treatment, as identified by our CAT method, can predict treatment response. This finding highlights a promising direction for biomarker development. A practical limitation, however, is that our current approach relies on single-cell RNA-seq data from tumor samples, which are often difficult to obtain. One potential solution is to detect CR T cells in blood, though enrichment strategies (e.g., enrichment of likely CR T cells by flow cytometry) will be necessary to reduce the cost of scRNA-seq.

Our association analysis can only identify TCRs for somatic mutations that appear in multiple individuals. Therefore, our NeoTCR platform is particularly suited for discovering public neoantigens shared across patients. Such public neoantigens provide opportunities for more cost-effective treatment options [22]. For example, vaccines could be developed as off-the-shelf products, in contrast to most personalized cancer vaccines, which are expensive and time-consuming to manufacture.

In addition, targeting public neoantigens identified by our NeoTCR platform may lead to more durable treatment responses. A major challenge in T cell–based therapies and vaccines is tumor immune evasion through down-regulation or loss of targeted neoantigens. Most neoantigens identified by our NeoTCR platform originate from public mutations, specifically from mutations in cancer driver genes that are essential for tumor survival. Thus, they are unlikely to be down-regulated or eliminated, reducing the risk of immune escape. For example, among the 47 somatic mutations prioritized as candidate neoantigen sources (Supplementary Table 5), eight have experimental evidence supporting their immunogenicity, and all eight arise from well-established oncogenes or tumor suppressor genes, including BRAF (V600E), KRAS (G12C, G12V), NRAS (Q61R, Q61K), PIK3CA (H1047R), and TP53 (R248Q, R248W) (Supplementary Table 6).

Our results further show that CR T cells from diverse patients and cancer types can be identified with high precision using gene expression data, whereas TCR sequence alone cannot reliably distinguish CR from non-CR T cells. This is likely because different cancer-reactive TCRs recognize distinct neo-peptide–HLA (npHLA) complexes and therefore do not share common sequence features. Nevertheless, this does not exclude the possibility that CR TCRs, particularly those restricted to a specific cancer type, may exhibit shared sequence motifs [16,17,36]. Moreover, recent advances in predicting TCR–antigen interactions directly from sequence data [37–40] could be integrated with our NeoTCR framework to further enhance the accuracy of detecting neoantigen-specific TCRs.

Our findings also suggest that TCR- or T cell–based features could serve as liquid biopsy biomarkers for cancer early detection. Current liquid biopsy approaches primarily rely on detecting epigenetic markers (e.g., DNA methylation or fragmentomics) from circulating tumor DNA (ctDNA), a subset of cell-free DNA (cfDNA). A key limitation is the low abundance of ctDNA, especially in early-stage cancers, compounded by the short half-life of cfDNA (5–150 minutes) [41]. In contrast, CR T cells can circulate for weeks to months, potentially making them easier to detect. Although circulating CR T cells are relatively rare (e.g., 1–5% as reported by [2]), they can be enriched using cell surface markers such as CD39 and CD103 [1, 5, 42] or by gene expression signatures, as shown in this study. Future studies may identify additional combinations of cell surface protein markers that further enhance the selective enrichment and detection of CR T cells.

Finally, our work points to several directions for improvement. First, the training data used to derive TCR–mutation associations were based on bulk RNA-seq with very limited TCR coverage and without paired alpha and beta chain information. We anticipate that datasets generated using cost-effective methods with paired chains and high read depth will substantially improve mutation prediction accuracy. Second, our analysis of TCR–mutation associations did not account for HLA restriction due to limited representation of public TCRs across populations. With larger datasets and higher TCR coverage, incorporating HLA information should further improve prediction accuracy, particularly for individuals with rare HLA alleles.

## Methods

### CR T gene signatures

In this paper, a gene signature refers to a set of genes, which may represent either a positive signature (genes with higher expression in cancer-reactive (CR) T cells) or a negative signature (genes with lower expression in CR T cells or higher expression in virus-specific T cells). Signatures were identified separately for CD8+ and CD4+ T cells. For each signature, we scored it by applying single-sample gene set enrichment analysis (ssGSEA) when it contained more than 30 genes [43]; otherwise, we used the average expression across genes. Below, we provide a brief overview of the gene expression signatures used in this study, with the exact sources listed in Supplementary Table 1.

Oliveira et al. [2] identified CR TCRs by testing up-regulation of the CD137 protein in TCR-transduced T cells when co-cultured with patient-derived melanoma cell lines (pdMel-CL). They then identified a set of genes that were differentially expressed in CD8 + T cells that had those CR TCRs. We extracted gene expression from Table S6 of Oliveira et al., where 74 and 26 genes were labeled tumor-specific and virus-specific, respectively.

Both Hanada et al. [5] and Lowery et al. [4] used a tandem minigene/peptide screening platform to identify functional CD8+ and CD4+ neoantigen-specific tumor-infiltrating T cells (TILs). Hanada et al. studied TILs from non-small cell lung cancer and Lowery et al. studied TILs from metastasis of several types of solid tumors. We recorded the gene expression signatures of Hanada et al. based on the results shown in Figure 6(D) (CD8+ T cells, 27/5 genes for positive/negative signature, respectively) and Figure 6(G) (CD4+ T cells, 9/4 genes for positive/negative signature, respectively) of Hanada et al. [5].

Lowery et al. [4] reported a 243-gene CD8+ positive signature and a 40-gene CD4+ positive signature (Table S4 of Lowery et al. [4]). We also extracted a 99-gene CD8+ negative signature and a 37-gene CD4+ negative signature from the differentially expressed genes reported in Table S8 of Lowery et al., using the criteria of average log fold change *<* 0, and adjusted p-value *<* 0.05. Lowery et al. also evaluated many published gene signatures and reported the signatures and AUCs in Table S4 and S12 of their paper, respectively. We chose to a positive signature with high AUC for CD8+ T cells: Oliveira TTE 100g (100 genes of terminally exhausted T cells while “100g” indicating the number of genes of the signature), and two positive signatures for CD4+ T cells: Oh CXCL13 50g and Caushi Tfh2 66g.

### TCR-mutation association analysis using TCGA data

We downloaded the TCR data of more than 8,000 cancer patients reported by Thorsson et al. [18]. There were in total 392,197 records of TCRs. Some TCRs were not from tumor tissues or contained non-amino acid sequence such as “∼” or “*”. After removing such entries, we were left with 333,263 TCRs. Approximately 21% of them had a unique V gene, 31% had no V genes, and the remaining had more than one V gene. Since most TCRs did not have a unique V gene, we ignored the V gene and only used CDR3 sequences to define TCRs.

We downloaded the somatic mutation calls of 9,243 cancer patients from a pancancer study [44]. We kept the mutations that were called by more than one caller and had a FILTER field being “PASS”, “wga”, or “native wga mix”. Next, we selected non-silent mutations belonging to one of the following groups: “Nonstop Mutation”, “Translation Start Site”, “In Frame Ins”, “In Frame Del”, “Splice Site”, “Nonsense Mutation”, “Frame Shift Ins”, “Frame Shift Del”, and “Missense Mutation”.

By intersecting patients with TCR data and those with somatic mutation data, we identified a cohort of 8,463 patients. From this group, we selected TCRs and somatic mutations present in at least five patients. This selection resulted in 4,619 somatic mutations, 2,048 TCR alpha chains, and 867 TCR beta chains.

We evaluated the association between each TCR and somatic mutation using logistic regression, modeling the probability of TCR detection as a function of TCR read depth, somatic mutation status, and their interaction. The significance of each association was assessed with a two–degree-of-freedom likelihood ratio test. The interaction term was included to reflect the expectation that neoantigen-specific TCRs may show increased abundance with read depth only when the corresponding mutation is present. Overall, the results were broadly similar with or without the interaction term (Supplementary Figure 8).

The TCGA dataset spans multiple cancer types with varying levels of T cell infiltration (Supplementary Figure 9). To test whether TCR–mutation associations were confounded by cancer type, we included cancer type as a covariate and recalculated association p-values. For each TCR–mutation pair, the maximum p-value across cancer types was taken as the adjusted measure of association strength. These values closely matched those obtained without adjustment (Supplementary Figures 10), indicating that cancer type had little impact on the observed associations.

### Validation of TCGA mutation prediction models with TRACERx dataset

We downloaded the TCR-seq reads of 68 patients from the TRACERx study [19] and called TCRs using decombinator (v5.0.0.dev0) [45], yielding 1,045,382 TCRs in tumor samples with amino-acid sequence length ≥ 5. We also downloaded the processed somatic mutations of 421 patients from the study [46]. By intersecting patients with TCR data and those with somatic mutation data, we were left with 65 patients. From this group, we selected somatic mutations present in at least two patients, yielding 62 mutations.

Among the 62 mutations, 11 had TCGA-trained models. To test the prediction accuracy of each TCGA-trained model on the 65 TRACERx patients, we first computed a TCR score for each patient defined as the fraction of that patient’s total TCR counts assigned to the model-selected TCRs. We then used this score to predict the binary mutation label and calculated the ROC AUC. This was done for TCR alpha and beta chains separately.

### False discovery rate (FDR) estimates for AUCs

We estimated a false discovery rate (FDR) for each AUC threshold, leveraging the fact that under the null hypothesis of no predictive power, AUC values should be symmetric around 0.5 [47]. Specifically, for the *j*-th prediction with AUC *y_j_*, we defined the FDR at cutoff 0.5+*δ* as *r_j_* = [*y_j_ <* (0.5−*δ*)]*/* [*y_j_ >* (0.5 + *δ*)]. See Supplementary Figure 8 for an illustration of the the estimates of FDR for different AUC cutoffs.

### Neural networks to classify CR versus non-CR T cells

We used the neural network to aggregate genome-wide gene expression to predict CR status. The CR label for each CD8+ or CD4+ T cell was generated as previously described (Supplementary Figure 2). We trained neural networks for CD8+ T cells and CD4+ T cells separately, using the same neural network architecture: an auto-encoder with a classification output (Supplementary Figure 4). The goal is to build a uniform definition of CR status across all the five studies. Therefore, for each study, we trained a model using all other studies and made prediction for the study of interest. The prediction score is highly consistent with the CR label in the left-out cancer, with AUCs larger than 0.92 across all studies. The neural network prediction scores are used to further filter out CR cells as those with prediction score larger than 0.5.

## Data availability

TCGA TCR data were derived from file TCGA mitcr cdr3 result 161008.tsv from https://gdc.cancer.gov/about-data/publications/panimmune. TCGA somatic mutation data were obtained from file mc3.v0.2.8.CONTROLLED.maf.gz from https://gdc.cancer.gov/about-data/publications/pancanatlas. Single-cell RNA-seq and TCR sequencing datasets analyzed in this study were obtained from the Gene Expression Omnibus (GEO) accession numbers: GSE156728 (pan-cancer [10]), GSE179994 (lung cancer [11]), GSE212217 (endometrial cancer [12]), GSE236581 (colorectal cancer [13]) and GSE243013 (lung cancer [14]).

## Software availability

CAT is available at https://github.com/Sun-lab/CAT. Neo-TCR is available at https://github.com/Sun-lab/neo-TCR.

## Supporting information

Supplementary results

## References

[1] Simoni, Y., Becht, E., Fehlings, M., Loh, C. Y., Koo, S.-L., Teng, K. W. W., Yeong, J. P. S., Nahar, R., Zhang, T., Kared, H., et al. (2018) Bystander CD8+ T cells are abundant and phenotypically distinct in human tumour infiltrates. Nature, 557(7706), 575–579.

[2] Oliveira, G., Stromhaug, K., Klaeger, S., Kula, T., Frederick, D. T., Le, P. M., Forman, J., Huang, T., Li, S., Zhang, W., et al. (2021) Phenotype, specificity and avidity of antitumour CD8+ T cells in melanoma. Nature, 596(7870), 119–125.

[3] Caushi, J. X., Zhang, J., Ji, Z., Vaghasia, A., Zhang, B., Hsiue, E. H.-C., Mog, B. J., Hou, W., Justesen, S., Blosser, R., et al. (2021) Transcriptional programs of neoantigen-specific TIL in anti-PD-1-treated lung cancers. Nature, 596(7870), 126–132.

[4] Lowery, F. J., Krishna, S., Yossef, R., Parikh, N. B., Chatani, P. D., Zacharakis, N., Parkhurst, M. R., Levin, N., Sindiri, S., Sachs, A., et al. (2022) Molecular signatures of antitumor neoantigen-reactive T cells from metastatic human cancers. Science, 375(6583), 877–884.

[5] Hanada, K.-i., Zhao, C., Gil-Hoyos, R., Gartner, J. J., Chow-Parmer, C., Lowery, F. J., Krishna, S., Prickett, T. D., Kivitz, S., Parkhurst, M. R., et al. (2022) A phenotypic signature that identifies neoantigen-reactive T cells in fresh human lung cancers. Cancer Cell, 40(5), 479–493.

[6] Veatch, J. R., Lee, S. M., Shasha, C., Singhi, N., Szeto, J. L., Moshiri, A. S., Kim, T. S., Smythe, K., Kong, P., Fitzgibbon, M., et al. (2022) Neoantigen-specific CD4+ T cells in human melanoma have diverse differentiation states and correlate with CD8+ T cell, macrophage, and B cell function. Cancer cell, 40(4), 393–409.

[7] Mair, F., Erickson, J. R., Frutoso, M., Konecny, A. J., Greene, E., Voillet, V., Maurice, N. J., Rongvaux, A., Dixon, D., Barber, B., et al. (2022) Extricating human tumour immune alterations from tissue inflammation. Nature, 605(7911), 728–735.

[8] Yost, K. E., Satpathy, A. T., Wells, D. K., Qi, Y., Wang, C., Kageyama, R., McNamara, K. L., Granja, J. M., Sarin, K. Y., Brown, R. A., et al. (2019) Clonal replacement of tumor-specific T cells following PD-1 blockade. Nature medicine, 25(8), 1251–1259.

[9] Oh, D. Y., Kwek, S. S., Raju, S. S., Li, T., McCarthy, E., Chow, E., Aran, D., Ilano, A., Pai, C.-C. S., Rancan, C., et al. (2020) Intratumoral CD4+ T cells mediate anti-tumor cytotoxicity in human bladder cancer. Cell, 181(7), 1612–1625.

[10] Zheng, L., Qin, S., Si, W., Wang, A., Xing, B., Gao, R., Ren, X., Wang, L., Wu, X., Zhang, J., et al. (2021) Pan-cancer single-cell landscape of tumor-infiltrating T cells. Science, 374(6574), abe6474.

[11] Liu, B., Hu, X., Feng, K., Gao, R., Xue, Z., Zhang, S., Zhang, Y., Corse, E., Hu, Y., Han, W., et al. (2022) Temporal single-cell tracing reveals clonal revival and expansion of precursor exhausted T cells during anti-PD-1 therapy in lung cancer. Nature cancer, 3(1), 108–121.

[12] Chow, R. D., Michaels, T., Bellone, S., Hartwich, T. M., Bonazzoli, E., Iwasaki, A., Song, E., and Santin, A. D. (2023) Distinct mechanisms of mismatch-repair deficiency delineate two modes of response to anti–PD-1 immunotherapy in endometrial carcinoma. Cancer discovery, 13(2), 312– 331.

[13] Chen, Y., Wang, D., Li, Y., Qi, L., Si, W., Bo, Y., Chen, X., Ye, Z., Fan, H., Liu, B., et al. (2024) Spatiotemporal single-cell analysis decodes cellular dynamics underlying different responses to immunotherapy in colorectal cancer. Cancer Cell, 42(7), 1268–1285.

[14] Liu, Z., Yang, Z., Wu, J., Zhang, W., Sun, Y., Zhang, C., Bai, G., Yang, L., Fan, H., Chen, Y., et al. (2025) A single-cell atlas reveals immune heterogeneity in anti-PD-1-treated non-small cell lung cancer. Cell, 188(11), 3081–3096.

[15] Goncharov, M., Bagaev, D., Shcherbinin, D., Zvyagin, I., Bolotin, D., Thomas, P. G., Minervina, A. A., Pogorelyy, M. V., Ladell, K., McLaren, J. E., et al. (2022) VDJdb in the pandemic era: a compendium of T cell receptors specific for SARS-CoV-2. Nature methods, 19(9), 1017–1019.

[16] Dash, P., Fiore-Gartland, A. J., Hertz, T., Wang, G. C., Sharma, S., Souquette, A., Crawford, J. C., Clemens, E. B., Nguyen, T. H., Kedzierska, K., et al. (2017) Quantifiable predictive features define epitope-specific T cell receptor repertoires. Nature, 547(7661), 89–93.

[17] Glanville, J., Huang, H., Nau, A., Hatton, O., Wagar, L. E., Rubelt, F., Ji, X., Han, A., Krams, S. M., Pettus, C., et al. (2017) Identifying specificity groups in the T cell receptor repertoire. Nature, 547(7661), 94–98.

[18] Thorsson, V., Gibbs, D. L., Brown, S. D., Wolf, D., Bortone, D. S., Yang, T.-H. O., Porta-Pardo, E., Gao, G. F., Plaisier, C. L., Eddy, J. A., et al. (2018) The immune landscape of cancer. Immunity, 48(4), 812–830.

19. Joshi, K., de Massy, M. R., Ismail, M., Reading, J. L., Uddin, I., Woolston, A., Hatipoglu, E., Oakes, T., Rosenthal, R., Peacock, T., et al. (2019) Spatial heterogeneity of the T cell receptor repertoire reflects the mutational landscape in lung cancer. Nature medicine, 25(10), 1549–1559.

[20] Reynisson, B., Alvarez, B., Paul, S., Peters, B., and Nielsen, M. (2020) NetMHCpan-4.1 and NetMHCIIpan-4.0: improved predictions of MHC antigen presentation by concurrent motif deconvolution and integration of MS MHC eluted ligand data. Nucleic acids research, 48(W1), W449–W454.

[21] Zhou, L. Y., Zou, F., and Sun, W. (2023) Prioritizing candidate peptides for cancer vaccines through predicting peptide presentation by HLA-I proteins. Biometrics, 79(3), 2664–2676.

[22] Pearlman, A. H., Hwang, M. S., Konig, M. F., Hsiue, E. H.-C., Douglass, J., DiNapoli, S. R., Mog, B. J., Bettegowda, C., Pardoll, D. M., Gabelli, S. B., et al. (2021) Targeting public neoantigens for cancer immunotherapy. Nature cancer, 2(5), 487–497.

23. Turajlic, S., Litchfield, K., Xu, H., Rosenthal, R., McGranahan, N., Reading, J. L., Wong, Y. N. S., Rowan, A., Kanu, N., Al Bakir, M., et al. (2017) Insertion-and-deletion-derived tumour-specific neoantigens and the immunogenic phenotype: a pan-cancer analysis. The lancet oncology, 18(8), 1009–1021.

[24] Xia, J., Bai, P., Fan, W., Li, Q., Li, Y., Wang, D., Yin, L., and Zhou, Y. (2021) NEPdb: a database of T-cell experimentally-validated neoantigens and pan-cancer predicted neoepitopes for cancer immunotherapy. Frontiers in Immunology, 12, 644637.

[25] Tran, E., Ahmadzadeh, M., Lu, Y.-C., Gros, A., Turcotte, S., Robbins, P. F., Gartner, J. J., Zheng, Z., Li, Y. F., Ray, S., et al. (2015) Immunogenicity of somatic mutations in human gastrointestinal cancers. Science, 350(6266), 1387–1390.

[26] Deniger, D. C., Pasetto, A., Robbins, P. F., Gartner, J. J., Prickett, T. D., Paria, B. C., Malekzadeh, P., Jia, L., Yossef, R., Langhan, M. M., et al. (2018) T-cell responses to TP53 “hotspot” mutations and unique neoantigens expressed by human ovarian cancers. Clinical Cancer Research, 24(22), 5562–5573.

[27] Malekzadeh, P., Pasetto, A., Robbins, P. F., Parkhurst, M. R., Paria, B. C., Jia, L., Gartner, J. J., Hill, V., Yu, Z., Restifo, N. P., et al. (2021) Neoantigen screening identifies broad TP53 mutant immunogenicity in patients with epithelial cancers. The Journal of clinical investigation, 129(3).

[28] Peri, A., Greenstein, E., Alon, M., Pai, J. A., Dingjan, T., Reich-Zeliger, S., Barnea, E., Barbolin, C., Levy, R., Arnedo-Pac, C., et al. (2021) Combined presentation and immunogenicity analysis reveals a recurrent RAS. Q61K neoantigen in melanoma. The Journal of clinical investigation, 131(20).

[29] Chandran, S. S., Ma, J., Klatt, M. G., Dündar, F., Bandlamudi, C., Razavi, P., Wen, H. Y., Weigelt, B., Zumbo, P., Fu, S. N., et al. (2022) Immunogenicity and therapeutic targeting of a public neoantigen derived from mutated PIK3CA. Nature medicine, 28(5), 946–957.

[30] McShan, A. C., Flores-Solis, D., Sun, Y., Garfinkle, S. E., Toor, J. S., Young, M. C., and Sgourakis, N. G. (2023) Conformational plasticity of RAS Q61 family of neoepitopes results in distinct features for targeted recognition. Nature Communications, 14(1), 8204.

[31] Kim, S. P., Vale, N. R., Zacharakis, N., Krishna, S., Yu, Z., Gasmi, B., Gartner, J. J., Sindiri, S., Malekzadeh, P., Deniger, D. C., et al. (2022) Adoptive cellular therapy with autologous tumor-infiltrating lymphocytes and T-cell receptor–engineered T cells targeting common p53 neoantigens in human solid tumors. Cancer immunology research, 10(8), 932–946.

[32] Parkhurst, M., Goff, S. L., Lowery, F. J., Beyer, R. K., Halas, H., Robbins, P. F., Prickett, T. D., Gartner, J. J., Sindiri, S., Krishna, S., et al. (2024) Adoptive transfer of personalized neoantigen-reactive TCR-transduced T cells in metastatic colorectal cancer: phase 2 trial interim results. Nature Medicine, 30(9), 2586–2595.

[33] Emerson, R. O., DeWitt, W. S., Vignali, M., Gravley, J., Hu, J. K., Osborne, E. J., Desmarais, C., Klinger, M., Carlson, C. S., Hansen, J. A., et al. (2017) Immunosequencing identifies signatures of cytomegalovirus exposure history and HLA-mediated effects on the T cell repertoire. Nature genetics, 49(5), 659–665.

34. Snyder, T. M., Gittelman, R. M., Klinger, M., May, D. H., Osborne, E. J., Taniguchi, R., Zahid, H. J., Kaplan, I. M., Dines, J. N., Noakes, M. N., et al. (2020) Magnitude and dynamics of the T-cell response to SARS-CoV-2 infection at both individual and population levels. *medRxiv*,.

[35] Sewell, A. K. (2012) Why must T cells be cross-reactive?. Nature Reviews Immunology, 12(9), 669–677.

[36] Yu, X., Pan, M., Ye, J., Hathaway, C. A., Tworoger, S. S., Lea, J., and Li, B. (2024) Quantifiable TCR repertoire changes in prediagnostic blood specimens among patients with high-grade ovarian cancer. Cell Reports Medicine, 5(7).

[37] Lu, T., Zhang, Z., Zhu, J., Wang, Y., Jiang, P., Xiao, X., Bernatchez, C., Heymach, J. V., Gibbons, D. L., Wang, J., et al. (2021) Deep learning-based prediction of the T cell receptor–antigen binding specificity. Nature machine intelligence, 3(10), 864–875.

[38] Bradley, P. (2023) Structure-based prediction of T cell receptor: peptide-MHC interactions. elife, 12, e82813.

[39] Meynard-Piganeau, B., Feinauer, C., Weigt, M., Walczak, A. M., and Mora, T. (2024) TULIP: A transformer-based unsupervised language model for interacting peptides and T cell receptors that generalizes to unseen epitopes. Proceedings of the National Academy of Sciences, 121(24), e2316401121.

[40] Croce, G., Bobisse, S., Moreno, D. L., Schmidt, J., Guillame, P., Harari, A., and Gfeller, D. (2024) Deep learning predictions of TCR-epitope interactions reveal epitope-specific chains in dual alpha T cells. Nature Communications, 15(1), 3211.

[41] Song, P., Wu, L. R., Yan, Y. H., Zhang, J. X., Chu, T., Kwong, L. N., Patel, A. A., and Zhang, D. Y. (2022) Limitations and opportunities of technologies for the analysis of cell-free DNA in cancer diagnostics. Nature biomedical engineering, 6(3), 232–245.

42. Duhen, T., Duhen, R., Montler, R., Moses, J., Moudgil, T., De Miranda, N. F., Goodall, C. P., Blair, T. C., Fox, B. A., McDermott, J. E., et al. (2018) Co-expression of CD39 and CD103 identifies tumor-reactive CD8 T cells in human solid tumors. Nature communications, 9(1), 2724.

[43] Hänzelmann, S., Castelo, R., and Guinney, J. (2013) GSVA: gene set variation analysis for microarray and RNA-seq data. BMC bioinformatics, 14, 1–15.

[44] Ellrott, K., Bailey, M. H., Saksena, G., Covington, K. R., Kandoth, C., Stewart, C., Hess, J., Ma, S., Chiotti, K. E., McLellan, M., et al. (2018) Scalable open science approach for mutation calling of tumor exomes using multiple genomic pipelines. Cell systems, 6(3), 271–281.

[45] Peacock, T., Heather, J. M., Ronel, T., and Chain, B. (08, 2020) Decombinator V4: an improved AIRR-C compliant-software package for T-cell receptor sequence annotation?. Bioinformatics, 37(6), 876–878.

46. Frankell, A. M., Dietzen, M., Al Bakir, M., Lim, E. L., Karasaki, T., Ward, S., Veeriah, S., Colliver, E., Huebner, A., Bunkum, A., et al. (2023) The evolution of lung cancer and impact of subclonal selection in TRACERx. Nature, 616(7957), 525–533.

[47] Bamber, D. (1975) The area above the ordinal dominance graph and the area below the receiver operating characteristic graph. Journal of mathematical psychology, 12(4), 387–415.

