## Supplementary results for "Characterization of Cancer-Reactive T Cells and Neoantigen-specific T Cell Receptors"

[1]Biostatistics Program, Public Health Sciences Division, Fred Hutchinson Cancer Center, Seattle, Washington, USA
[2]Department of Statistics, University of Washington, Seattle, Washington, USA
[3]Herbold Computational Biology Program, Public Health Sciences Division, Fred Hutchinson Cancer Center, Seattle, Washington, USA
[4]Institute for Protein Design. University of Washington, Seattle, Washington, USA
[5]Department of Biostatistics, University of Washington, Seattle, Washington, USA
[6]Department of Biostatistics, University of North Carolina, Chapel Hill, North Carolina, USA
[7]Translational Science and Therapeutics Division, Fred Hutchinson Cancer Center, Seattle, WA, USA
[8]Seattle Children’s Hospital, Seattle, WA, USA
[9]Department of Pediatrics, University of Washington School of Medicine, Seattle, WA, USA

October 30, 2025

### Gene expression signatures for cancer reactive T cells

| Signature | Cell type | Direction | Source |
|---|---|---|---|
| Hanada_neg_4g | CD4 | negative | Hanada et al. Figure 6 |
| Lowery_neg_37g | CD4 | negative | Lowery et al. Table S8 |
| Caushi_Tfh2_66g | CD4 | positive | Lowery et al. Table S4 |
| Hanada_pos_9g | CD4 | positive | Hanada et al. Figure 6 |
| Lowery_pos_40g | CD4 | positive | Lowery et al. Table S4 |
| Oh_CXCL13_50g | CD4 | positive | Lowery et al. Table S4 |
| Hanada_neg_5g | CD8 | negative | Hanada et al. Figure 6 |
| Oliveira_virus_26g | CD8 | negative | Oliveira et al. Table S6 |
| Hanada_pos_27g | CD8 | positive | Hanada et al. Figure 6 |
| Lowery_pos_243g | CD8 | positive | Lowery et al. Table S4 |
| Oliveira_pos_74g | CD8 | positive | Oliveira et al. Table S6 |
| Oliveira_TTE_100g | CD8 | positive | Lowery et al. Table S4 |
| Yost_CD8_Exh_100g | CD8 | positive | Lowery et al. Table S4 |

Supplementary Table 1: List of gene signatures used in this work to identify cancer reactive T cells and non-cancer reactive T cells.

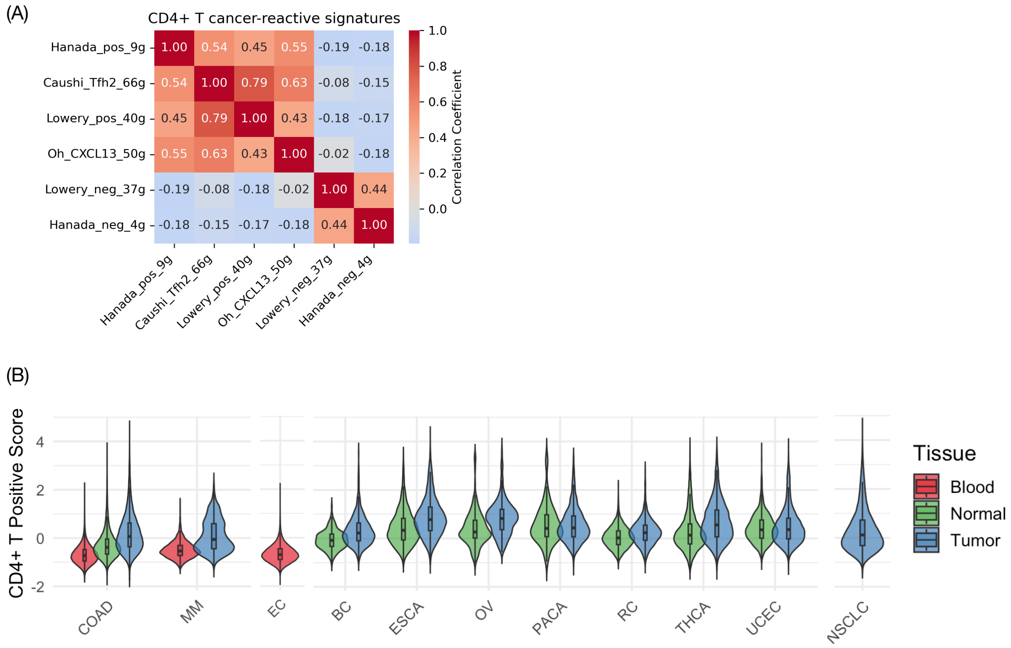


Supplementary Figure 1: (A) The correlation of different signatures. (B) The CD4+ positive signature scores across cancer types and tissue types.

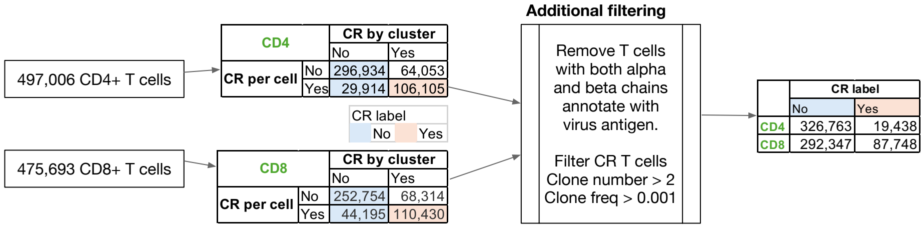


Supplementary Figure 2: The work flow to identify potential cancer-reactive T cells versus non-cancer-reactive T cells.

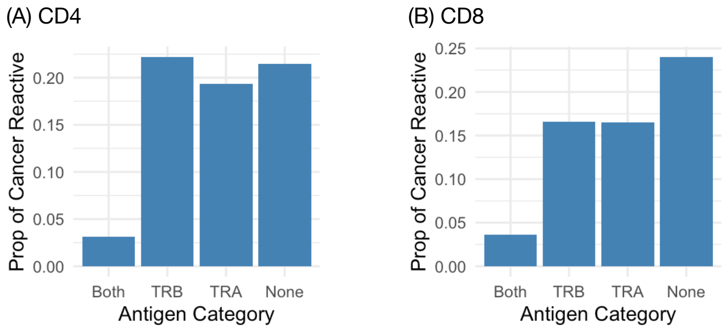


Supplementary Figure 3: The proportion of CD4+ T cells (A) or CD8+ T cells (B) classified as cancer reactive, stratified by wether both TCR alpha and beta chains are annotated with non-human antigens or just one of the two chains, or none of the two chains. Based on these results, we exclude those T cells, of which both alpha and beta chains are annotated with non-human antigens.

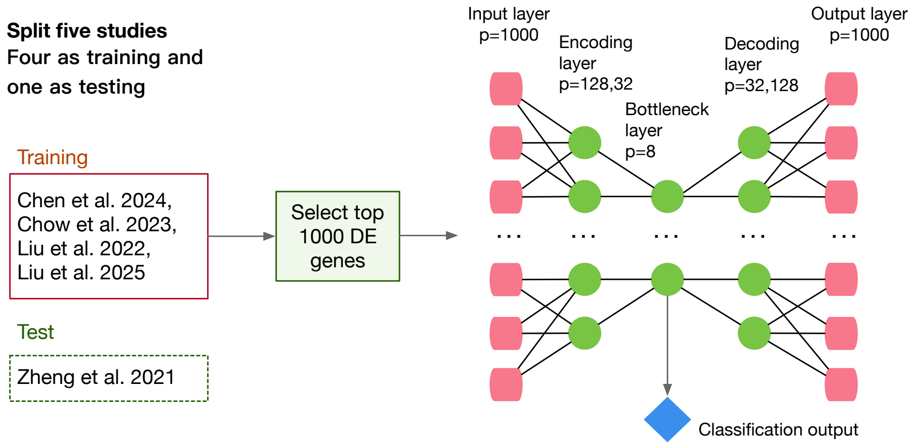


Supplementary Figure 4: Predict cancer reactive status of each T cell from one study by the neural network trained using the data from four other studies.

### CR T cells cannot be accurately identified by their TCRs

To avoid potential batch effects across studies and to select both CR and non-CR T cells, we focused on the T cells from one pan-cancer T cell study [1]. We clustered T cells using the expression of genes involved in CR signatures, and manually select cell clusters that are CR or non-CR and ignored those ambiguous clusters. For each cancer type, we clustered CD8+ and CD4+ T cells separately and calculated the median positive/negative scores for each cluster. We labeled a cluster as cancer-reactive if it had high median positive score and a low median negative score. Conversely, a cluster was labeled as non-cancer-reactive if it had a high median negative score and a low median positive score (Supplementary Figure 5). Through manual examination of the cluster-level scores, we identified cancer-reactive and non-cancer-reactive CD4+ and CD8+ T cells for each cancer type.

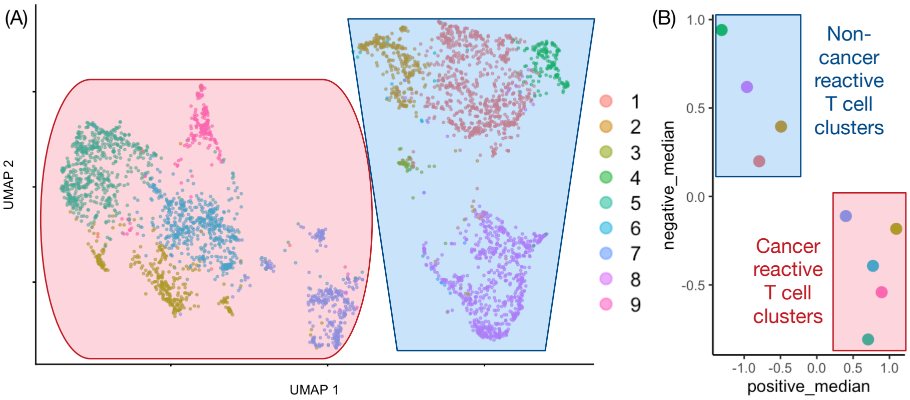


Supplementary Figure 5: Selection of cancer-reactive and non-cancer-reactive CD8+ T cells from Esophageal carcinoma. **(A)** UMAP projections with labels for nine clusters. **(B)** The median values of cancer-reactive (positive) and non-cancer-reactive (negative) scores for each cluster. The clusters selected as cancer-reactive or non-cancer-reactive CD8+ T cells are highlighted.

After combining the annotated TCRs across cancer types, we identified 6,367 cancer-reactive TCRs and 15,685 non-cancer-reactive TCRs. We classified them using multiple neural networks with different input encodings and neural network architectures. Specifically, neural networks were constructed according to four factors: input feature, encoding method, dense layer size, and # of CNN layers (Supplementary Table 2). This resulted in $6 \times 4 \times 4 \times 2 = 192$ neural network models. The output of each neural network was a number between 0 and 1 indicating the confidence level that a TCR was CR. Supplementary Figure 6(A) illustrates one neural network. The evaluation for each neural network model was done based on average prediction scores for each test TCR given by an ensemble of 20 models trained on 20 different random seeds.

| **Factor** | **Options** |
|---|---|
| Input Feature | (alpha chain only, beta chain only, both alpha and beta chain) × (CDR3 only, CDR3 + V gene) |
| Encoding Method | one hot, blosum62, Atchley, and PCA |
| Dense Layer Size | (32, 16), (64, 32), (128, 64), (256, 128) |
| # of CNN Layers | 1, 2 |

Supplementary Table 2: Different options for the neural network models to classify cancer reactive TCRs versus non-cancer reactive TCRs.

The rationale for considering many neural network models was to evaluate the feasibility of predicting CR TCRs from their sequences, regardless of the optimal model. The AUCs were higher when using the beta chain alone compared to the alpha chain alone, possibly due to the higher complexity of the beta chains (Figure 6(B)). On average, using both alpha and beta chains, along with V genes, resulted in the highest AUCs. However, even the highest AUC was below 0.6 (Figure 6(B)). In other words, the probability that a CR TCR is ranked higher than a non-CR TCR was below 0.6. Therefore, we conclude that using this dataset, the signals in the TCRs are not strong enough to distinguish CR T cells and non-CR T cells.

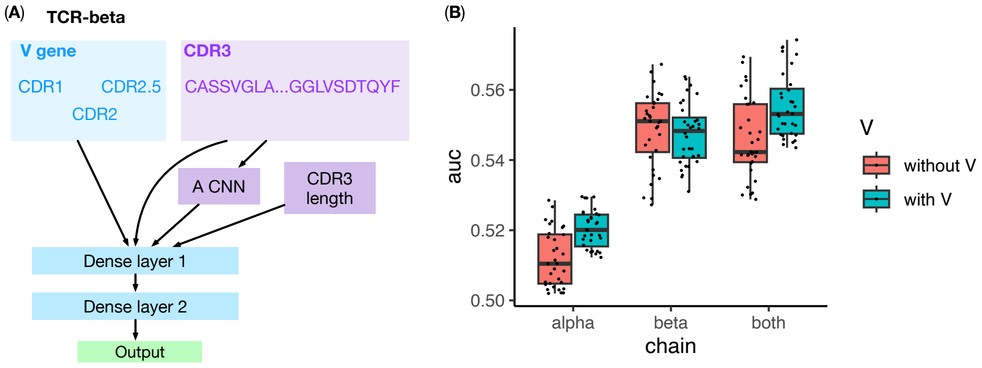


Supplementary Figure 6: **(A)** An example of the architecture of a neural network to classify CR TCRs versus non-CR TCRs. This neural network uses both the V gene and the CDR3 region of a TCR beta chain. A convolutional neural network (CNN) is applied to the CDR3 to capture position-independent features. The output of this CNN is then combined with the encoded CDR3 sequence as input for the classification task. **(B)** The distribution of AUCs in the testing data for CR TCR classification, using different input data and neural network hyper-parameters.

### Additional results for the analysis of TCGA somatic mutation and TCR data

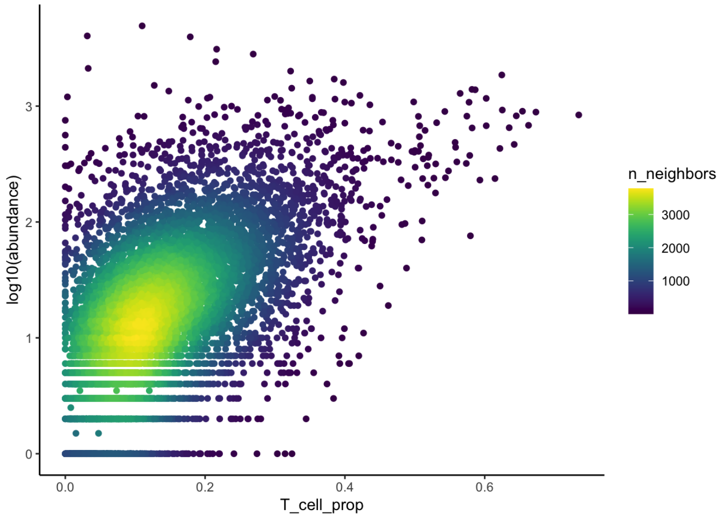


Supplementary Figure 7: The number of TCRs from each sample is associated with T cell proportion of each sample.

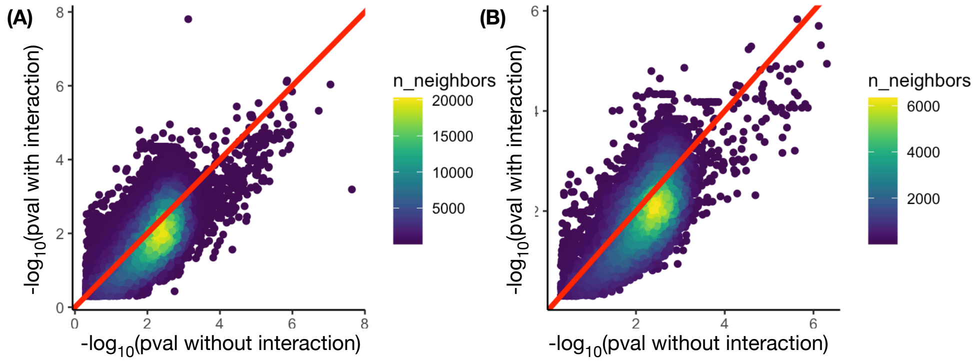


Supplementary Figure 8: Comparison of the p-values for TCR-mutation associations with or without interactions between mutations and TCR read-depth. (A) TCR alpha chain. (B) TCR beta chain.

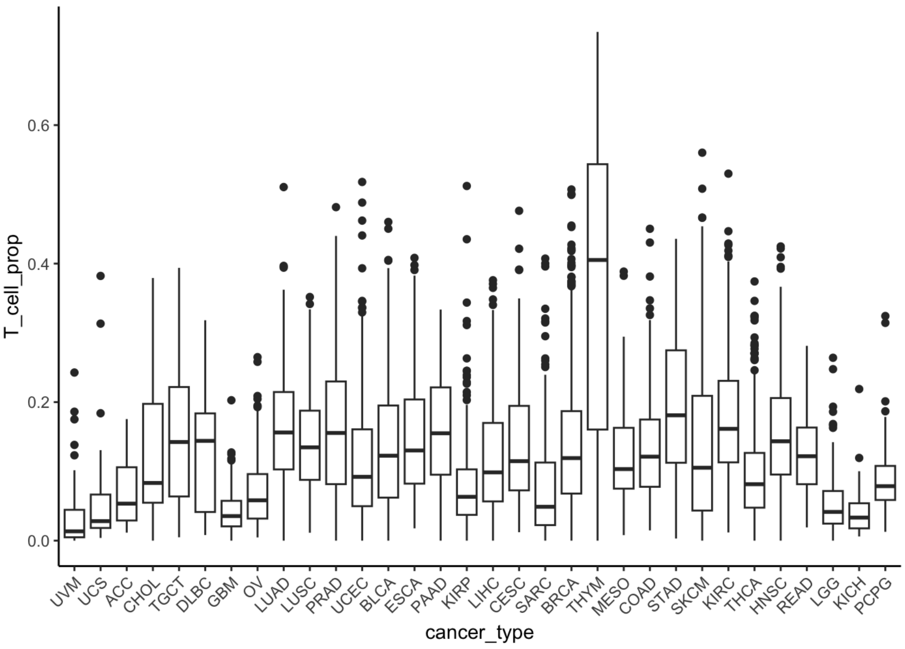


Supplementary Figure 9: Variation of T cell proportions across cancer types.

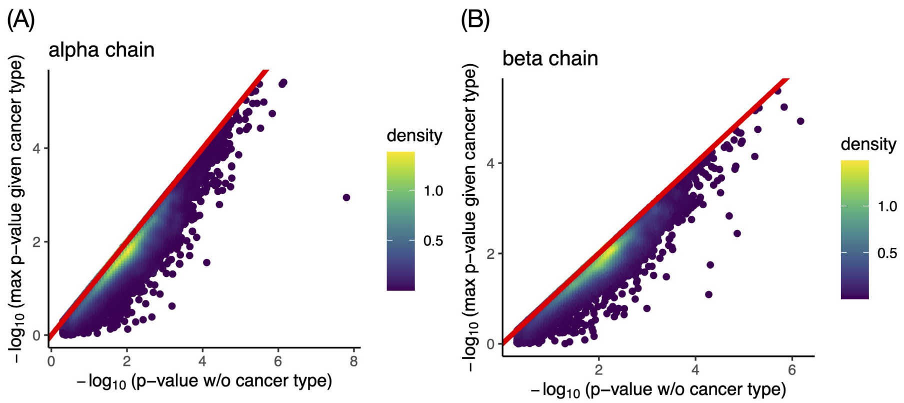


Supplementary Figure 10: Comparison of the p-values for TCR-mutation associations without accounting for cancer types versus the maximum of the p-values for TCR-mutation associations when accounting for each cancer type, for TCR alpha chain (A) and beta chain (B). Each point is a pair of somatic mutation and a TCR alpha or beta chain.

### Estimation of FDR for different AUC thresholds.

Since the alpha chain has less diversity, it is more likely that two individuals share the same TCR chain chain. Therefore, it is easier for alpha chain predictor to achieve high AUC by chance, and as shown in the following two figures, more stringent AUC cutoffs are needed for TCR alpha chain to control FDR.

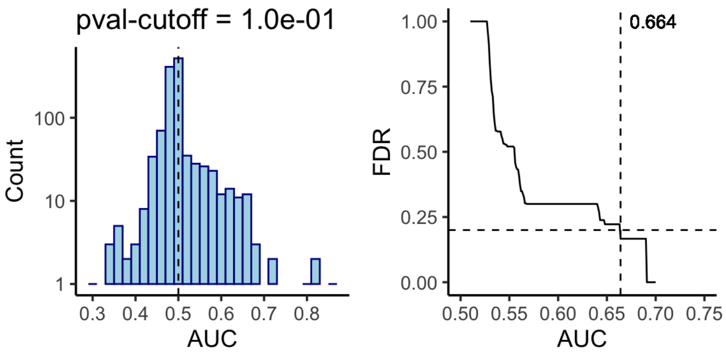


Supplementary Figure 11: Left panel: the distribution of AUCs for prediction by TCR **alpha** chain, when the p-value cutoff is 0.1. Right panel: the corresponding FDR for different AUC cutoffs.

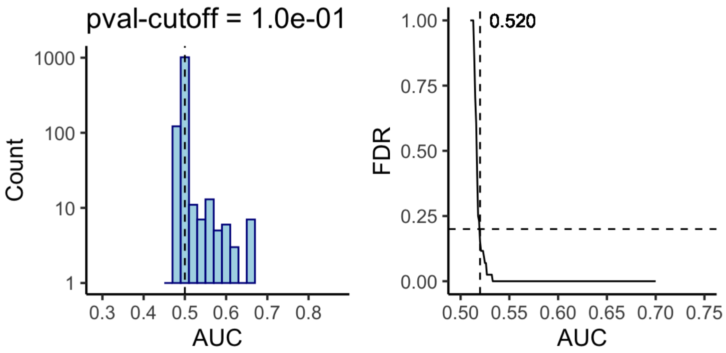


Supplementary Figure 12: Left panel: the distribution of AUCs for prediction by TCR **beta** chain, when the p-value cutoff is 0.1. Right panel: the corresponding FDR for different AUC cutoffs.

### Validation of TCR-based prediction.

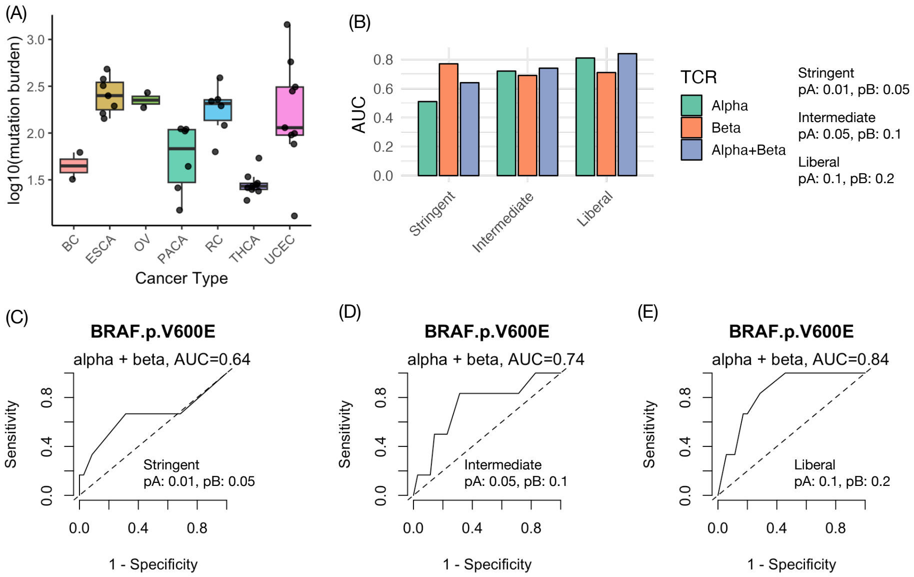


Supplementary Figure 13: Validation of TCR-based predictions of BRAF V600E mutation using the data of 41 patients reported by Zheng et al. [1]. (A) Summary of mutation burdens of 41 patients across multiple cancer types. (B) AUCs for three prediction models when selecting mutation-associated TCRs at three different p-value cutoffs (Stringent, Intermediate, and Liberal), and using TCR alpha chain only, beta chain only, or both alpha and beta chains. (C)-(E). ROC curves for three prediction models of BARF V600E mutation, while using both TCR alpha and beta chains.

### T cells with mutation-associated TCRs tend to have CR gene expression signatures

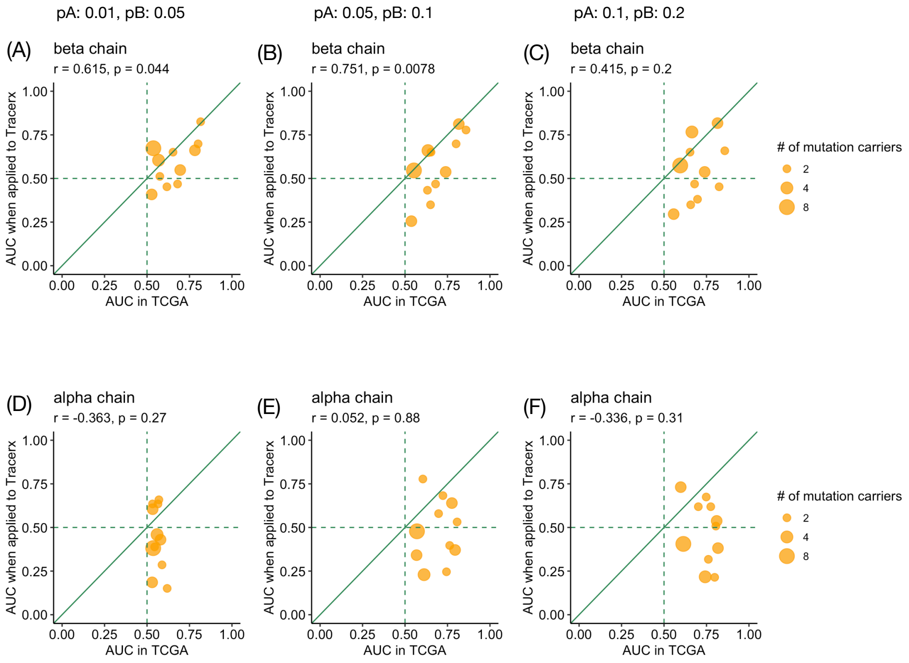


Supplementary Figure 14: Comparison of prediction performance between TCGA and TRACERx datasets. Shown are the AUCs for somatic mutation prediction in TCGA compared with the AUCs obtained when applying the same models to TRACERx data [2]. Three sets of models were constructed using different p-value thresholds for mutation–TCR associations, with cutoffs for the TCR alpha- and beta-chains denoted by pA and pB, respectively: (A, D) pA = 0.01, pB = 0.05; (B, E) pA = 0.05, pB = 0.1; and (C, F) pA = 0.1, pB = 0.2. Predictions based on the beta-chain (A–C) show strong correlation between TCGA and TRACERx, whereas predictions based on the alpha-chain (D–F) show no concordance.

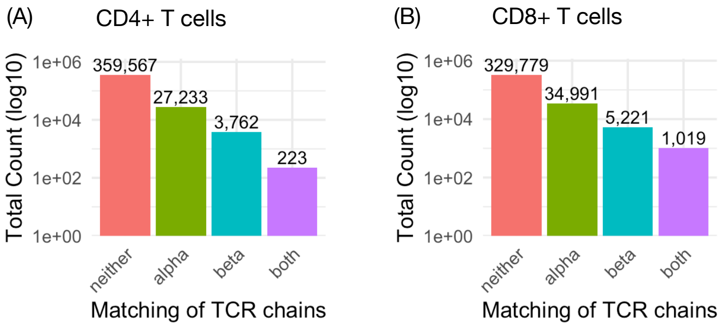


Supplementary Figure 15: The number of CD8+ T cells or CD4+ T cells classified based on whether their TCRs are associated with a somatic mutation from the TCGA analysis

### Characterization of different groups of somatic mutations

Compare different groups of somatic mutations stratified by their associations with TCRs: "No" association, "Weak" association or "Strong" association, related with Figure 5 of main text.

| Mutation Type | No | Strong | Weak |
|---|---|---|---|
| Frame_Shift_Del | 180 / 328 | 508 / 543 | 688 / 505 |
| Frame_Shift_Ins | 20 / 21 | 35 / 35 | 33 / 32 |
| In_Frame_Del | 5 / 6 | 14 / 11 | 8 / 10 |
| Missense_Mutation | 879 / 740 | 1258 / 1225 | 966 / 1138 |
| Nonsense_Mutation | 144 / 133 | 219 / 220 | 194 / 204 |

Supplementary Table 3: Observed vs Expected (Observed / Expected) number of mutations for each mutation type (each row) and each group in terms of associations with TCRs.

| | All | | Strong | | | | |
|---|---|---|---|---|---|---|---|
| **AA** | **Freq** | **Prop** | **Freq** | **Prop** | **Ratio** | **pvalue** | **odds ratio** |
| R → Q | 52956 | 0.024 | 237 | 0.19 | 8.0 | 2.2e-133 | 9.5 |
| S → L | 29231 | 0.013 | 103 | 0.08 | 6.3 | 1.1e-47 | 6.7 |
| R → C | 49135 | 0.022 | 150 | 0.12 | 5.5 | 1.3e-61 | 6.0 |
| R → I | 17162 | 0.0076 | 50 | 0.04 | 5.2 | 2.9e-20 | 5.3 |
| E → K | 83559 | 0.037 | 227 | 0.18 | 4.9 | 5.8e-85 | 5.6 |
| R → W | 37709 | 0.017 | 53 | 0.04 | 2.5 | 4.3e-09 | 2.6 |

Supplementary Table 4: The types of mis-sense mutations that are much more likely to be **public** neoantgen than any non-synonymous somatic mutations.

Supplementary Table 5: 47 somatic mutations prioritized as candidate neoantigen sources. See the excel file `sTable_5_mut_neo.xlsx`.

Supplementary Table 6: Summary of eight somatic mutations with experimental evidence for immunogenicity. The complete list of known neoantigens with relative references is available at `https://github.com/Sun-lab/neo-TCR/blob/main/somatic_mutation/data/konwn_public_neoantigen.xlsx`.

| **key_pro** | **key** | **cancer** | **auc** | **nTCRA** | **nTCRB** | **Freq** |
|---|---|---|---|---|---|---|
| BRAF:V600E | BRAF:7:140453136:A:T | | | 117 | 81 | 558 |
| PIK3CA:H1047R | PIK3CA:3:178952085:A:G | | | 64 | 47 | 242 |
| KRAS:G12V | KRAS:12:25398284:C:A | STAD | 0.641 | 72 | 48 | 169 |
| TP53:R248Q | TP53:17:7577538:C:T | LUSC | 0.852 | 54 | 36 | 121 |
| NRAS:Q61R | NRAS:1:115256529:T:C | | | 79 | 32 | 112 |
| KRAS:G12C | KRAS:12:25398285:C:A | | | 98 | 46 | 96 |
| TP53:R248W | TP53:17:7577539:G:A | STAD,OV | 0.829,0.633 | 55 | 36 | 89 |
| NRAS:Q61K | NRAS:1:115256530:G:T | | | 66 | 38 | 67 |

### References


[1] Zheng, L., Qin, S., Si, W., Wang, A., Xing, B., Gao, R., Ren, X., Wang, L., Wu, X., Zhang, J., et al. (2021) Pan-cancer single-cell landscape of tumor-infiltrating T cells. *Science,* **374**(6574), abe6474.

[2] Joshi, K., de Massy, M. R., Ismail, M., Reading, J. L., Uddin, I., Woolston, A., Hatipoglu, E., Oakes, T., Rosenthal, R., Peacock, T., et al. (2019) Spatial heterogeneity of the T cell receptor repertoire reflects the mutational landscape in lung cancer. *Nature medicine,* **25**(10), 1549–1559.
